# Loss of *dmrt1* restores female fates in the absence of *cyp19a1a* but not *rbpms2*

**DOI:** 10.1101/2020.03.26.009522

**Authors:** Shannon Romano, Odelya H. Kaufman, Florence L. Marlow

## Abstract

Sex determination and differentiation is a complex process regulated by multiple factors, including factors from the germline or surrounding somatic tissue. In zebrafish, sex-determination begins with establishment of a bipotential gonad that undergoes sex-specific differentiation and maintenance to form the functional adult gonad. However, the relationships among these factors are not fully understood. Here we identify potential Rbpms2 targets and apply genetic epistasis experiments to decipher the genetic hierarchy of regulators of sex-specific differentiation. We provide evidence that the critical female factor, *rbpms2* is epistatic to the male factor *dmrt1* in terms of adult sex. Moreover, Rbpms2’s role in promoting female fates extends beyond repression of Dmrt1, as Rbpms2 is essential for female differentiation even in the absence of Dmrt1. In contrast, female fates can be restored in mutants lacking *cyp19a1a* in the absence of *dmrt1.* Taken together this work indicates that Cyp19a1a-mediated suppression of Dmrt1 is key to establish a bipotential gonad and initiate female fate acquisition, possibly by promoting *rbpms2*. Then, after female fate specification, Cyp19a1a regulates subsequent oocyte maturation and sustains female fates independent of Dmrt1 repression.

**Author Summary:** We show that *cyp19a1a*-mediated suppression of *dmrt1* establishes a bipotential gonad and female fate acquisition, possibly through *rbpms2* which is required for female fates, even in the absence of Dmrt1.

## Introduction

Although sex-determination is a common biological process critical to reproduction and thus species survival in sexually reproducing organisms, the mechanisms controlling this process vary extensively among animal species. The decision to develop as one sex or the other is triggered in different species by many environmental factors including temperature and nutrition status, presence of sex chromosomes, and the function of many sex-associated genetic factors that converge upon regulation of downstream sexual differentiation factors, for example, *Sox9* (*SRY-box 9), FoxL2* (*Forkhead box L2), Dmrt1 (Doublesex and mab-3 related transcription factor 1)*, and *Cyp19a1a (Aromatase; Cytochrome P450, family 19, subfamily a*)[1–4]. In domesticated lab strains of zebrafish, sex determination appears to be polygenic[5–8], being influenced by numerous genetic factors. Like most teleost fish, zebrafish are a gonochoristic species, eventually differentiating into one of two sexes, that do not switch sex as adults[9]. However, the early zebrafish gonad is bipotential and thus is poised to develop as either an ovary or a testis[10]. This plasticity provides an excellent model for studying genetic factors and interactions between essential regulators of sex-specific differentiation among vertebrates.

Factors regulating sex differentiation and maintenance vary in function, from transcription factors regulating sex-specific programs, to regulators of signaling molecules, and even signaling molecules themselves. *Doublesex and mab-3 related transcription factor 1* (*Dmrt1*) is a transcription factor critical for male gonad differentiation and maintenance across vertebrates[2, 11–15]. In mammals, Dmrt1 acts by directly activating male determining genes and simultaneously suppressing female determining genes[2, 16]. Recent studies verified that *dmrt1* is expressed in both the germline and somatic gonad in zebrafish and as in mammals, is required for male differentiation, as mutants disrupting Dmrt1 cause female-biased sex ratios[11, 17]. Aromatase, encoded by *cyp19*, required for the final steps of estrogen synthesis[18, 19], is a critical factor for sex differentiation in fish[10, 20]. Zebrafish have two copies of *cyp19 (*ovarian *cyp19a1a* and brain *cyp19a1b)* and, in contrast to *dmrt1*, knockout of the ovarian aromatase *cyp19a1a* results in an all-male phenotype that is characterized by early transition to male gonad features with an apparent absence of a bipotential phase of development[21].

Bone morphogenetic protein 15 (Bmp15) is a signaling molecule expressed primarily by the oocyte in mice and zebrafish[20, 22, 23]. Bmp15 has recently been shown to be required in zebrafish for maintenance of the female gonad, after sex has been determined[20]. Knockout of *bmp15* in zebrafish causes early switching of mutant sex from female to male, and thus results in recovery of only fertile adult males[20]. Although work in mammalian systems indicates that TGF-β signaling, particularly GDF-9 and BMP15 are crucial for granulosa cell fate/development[24–26], GDF-9 appears to be dispensable in zebrafish as loss of function causes no overt phenotypes and does not worsen phenotypes caused by loss of *bmp15*[20]. Recent work from our lab identified an RNA binding protein, RNA-binding protein of multiple splice forms 2 (Rbpms2), as a critical germline expressed[27] factor for female sex differentiation in zebrafish[27]. Simultaneous loss of both redundant copies of *rbpms2* (*rbpms2a* and *rbpms2b; rbpms*2 double mutants (DMs)) results in recovery of exclusively fertile males[27]; however, how Rbpms2 promotes female fates is not clear. It has also been shown that both the somatic gonad and meiotic oocytes contribute to establishing and maintaining the female gonad in domesticated zebrafish (reviewed by [1]). Therefore, it is expected that the sex-specific identities of the somatic gonad and germ cells must match to successfully develop a functional gonad.

Although critical roles for each of the above-mentioned genes in determining sexual fate in zebrafish have been established, and it is known that female and male pathways function antagonistically[1, 28], precisely where each factor resides in the hierarchy of genes regulating germ cell and somatic gonad sex and their interactions in gonadal differentiation remains incompletely understood. In this work, we surveyed known regulators of sex determination to identify potential Rbpms2 targets. Specifically, we used genetic epistasis experiments and cell biological approaches to tease apart the genetic hierarchy of these critical factors in sex determination. We provide evidence that TGF-β signaling is activated in early germ cells of the bipotential gonad and in wild-type becomes activated in the somatic gonad upon sexual differentiation, and that in *rbpms2DM*s initial activation of TGF-β in the bipotential germline is intact. Since zebrafish establish a bipotential gonad that is initially female in character, we reasoned that antagonism of male promoting fates, such as Dmrt1 would be key to female-specific differentiation.

Surprisingly, we found that loss of *dmrt1* was not sufficient to suppress the male only differentiation phenotype of the *rbpms2DM*s, thus *rbpms2* is epistatic to *dmrt1* in terms of adult sex. Moreover, Rbpms2 has a role beyond simply repressing or antagonizing Dmrt1, as it is required to promote female fates and for female sex-specific differentiation even in the absence of Dmrt1. Interestingly, unlike either *rbpms2DM*s, which develop as fertile males, or *dmrt1*^*uc27*^ mutants, which develop as fertile females, *rbpms2;dmrt1TMs* were sterile in breeding and immunohistological assays, indicating that germ cells may be lost due to cell autonomous failure to establish sex-specific identity, nonautonomous defects in somatic gonad development, or as a consequence of mismatched germ cell and somatic gonad sex-specific identity. Further demonstrating the distinct contribution of Rbpms2 to promoting female fates, we found that loss of *dmrt1* was sufficient to suppress the early male-differentiating phenotype of *cyp19a1a* mutant gonads as evidenced by the presence of oocytes containing the female specific marker Buckyball and morphologically normal Balbiani bodies in *cyp19a1a;dmrt1DM* gonads [29]. In contrast, early *cyp19a1a* single mutant gonads prematurely take on a male identity and fail to develop any oocytes. Taken together these findings indicate that *cyp19a1a* acts during at least two steps of female specific differentiation. First, *cyp19a1a* mediated suppression of *dmrt1* is key to establish a bipotential gonad and initiate female fate acquisition in zebrafish, possibly by promoting *rbpms2* which is required for female specific differentiation, even in the absence of Dmrt1. Ultimately, female fates are not maintained in *cyp19a1a;dmrt1DMs* likely due to the later Bmp15-dependent expression of Cyp19a1a that is required for subsequent follicle differentiation and oocyte maintenance by a mechanism that is independent of inhibition of Dmrt1.

## Results and Discussion

### Molecular characterization of rbpms2DM juvenile gonads

We previously showed that cellular features of *rbpms2DM* juvenile gonads are comparable to wild-type[27]. To determine if *rbpms2DM* juvenile gonads resemble wild-type molecularly, we prepared cDNA from day 21 post fertilization (d21) trunks, prior to sex-determination, and used RTPCR to examine markers associated with female or male-specific somatic gonad or germ cell fates. Specifically, we examined *vasa* (all germ cells), *sox9a* (oocyte)[30], *cyp19a1a* (Granulosa cells)[20, 30], *cyp11c1* (Leydig cells), *amh* (Sertoli cells)[30], *bmpr1ab, bmpr1bb, bmpr2b*[20], and *ef1alpha* (control). We saw no differences in expression of these markers between *rbpms2* DMs and wild-type siblings at d21 (Fig. 1A), confirming that expression of *cyp19a1a* and the bipotential gonad are established in the absence of Rbpms2.

**Figure 1.**
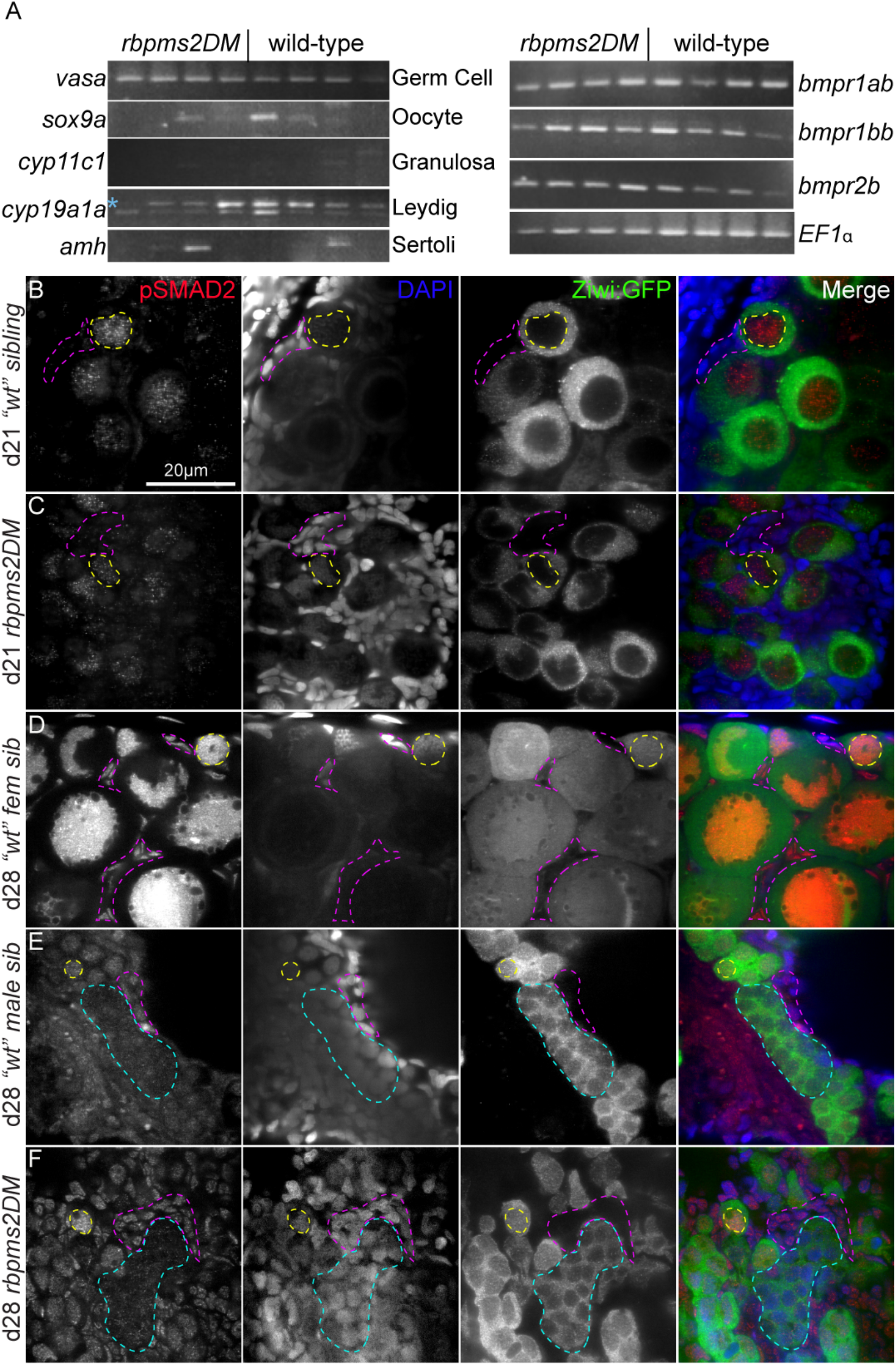
Molecularly bipotential gonads of *rbpms2DMs* and activation of p-SMAD in early germ cells and differentiating gonads. (A) RT-PCR on dissected trunks from d21 rbpms2 DMs and wildtype siblings reveal no differences in expression of relevant molecular markers (left panel, left column) for indicated cell type (left panel, right column) or for relevant receptors (right panel, right column).(B-F) p-SMAD2, DAPI and ziwi:GFP staining. Representative slices from Z-series reveal localization of p-SMAD2 in bipotential and early differentiating gonads of d21 and d28 juvenile zebrafish, respectively. In undifferentiated bipotential gonads of both (A) wild-type (n=3) and (B) *rbpms2DM* (n=3) p-Smad2 is detected in the nuclei of germ cells (example outlined with yellow dashed line), but not the nuclei of the surrounding somatic cells (examples outlined in pink dashed lines). By d28 in early differentiated (C) females (n=4), p-SMAD2 remains localized to the germ cell nuclei and begins to be expressed in somatic nuclei. Similarly, p-SMAD2 is expressed at d28 in both germ cell nuclei (examples outlined in teal dashed lines) and somatic nuclei of (D) wild-type male (n=4) and (E) *rbpms2DM* gonads (n=4).

### TGF-β signaling in the differentiating gonad

In mammalian systems, TGF-β signaling mediated by GDF-9 and BMP15 form an oocyte-granulosa cell feedback loop[25, 26, 31–35]; however, in zebrafish GDF-9 appears to be dispensable because *gdf-9* loss neither causes overt phenotypes nor worsens phenotypes caused by loss of *bmp15*[20]; thus, the role of TGF-β in the differentiating zebrafish gonad remains unclear. To investigate potential involvement of TGF-β, we examined the localization of p-SMAD2, an indicator of active TGF-β signaling[36–38]. In undifferentiated (d21) wild-type (Figure 1B) and *rbpms2DM* (Figure 1C) gonads, p-SMAD2 was detected with DAPI in the nuclei of germ cells marked by ziwi-GFP[39]. As differentiation proceeded in wild-type females, p-SMAD2 remained strong in the nuclei of early oocytes and became activated in the nuclei of somatic gonad cells (Figure 1D). Nuclear p-SMAD2 was also detected in the somatic gonad of wild-type (Fig. 1E) and *rbpms2DM* (Fig. 1F) males, albeit appeared weaker than in females and declines in differentiating spermatogonia. Based on these observations, we conclude that TGF-β is active in the early germline and soma of the differentiating gonads of wild-type, reminiscent of the patterns observed in mammals[40]; however, the relevant ligand(s) in zebrafish remains to be identified. Moreover, although initial activation of p-SMAD2 in the germline was intact, the strong signals achieved in germline and soma of wild-type females were not detected in *rbpms2DM*s. Because Rbpms2 binds to and has been implicated in potentiating activation of Smad2/3 targets[41], it is possible that Rpbms2 promotes female fates by boosting TGF-β signaling; however, it remains to be determined if this is a cause or effect of failed female-specific differentiation.

### Identification of candidate Rbpms2 target RNAs

Because *rbpms2* double mutants can develop as fertile males that complete meiosis and spermatogenesis but fail to complete female differentiation[27], we reasoned that *rbpms2* is not required for meiosis per se, but instead is required to prevent the germline from adopting a male fate. This could occur either by promoting expression of female factors, by repressing male factors, or a combination of both. Since Rbpms2 is an RNA-binding protein, we surveyed genes required for meiosis, oogenesis, and sex determination for presence of the previously identified Rbpms RNA-binding site consensus sequence, *CAC(n3-12)CAC* [42]. We identified five or more Rbpms RNA-binding sites within the untranslated regions (*utrs)* and exons of the male fate regulator, *dmrt1*, the TGF-β signaling molecule, *bmp15*, and aromatase, *cyp19a1a (*Table 1), making them compelling targets gene for regulation by Rbpms2.

**Table 1.**
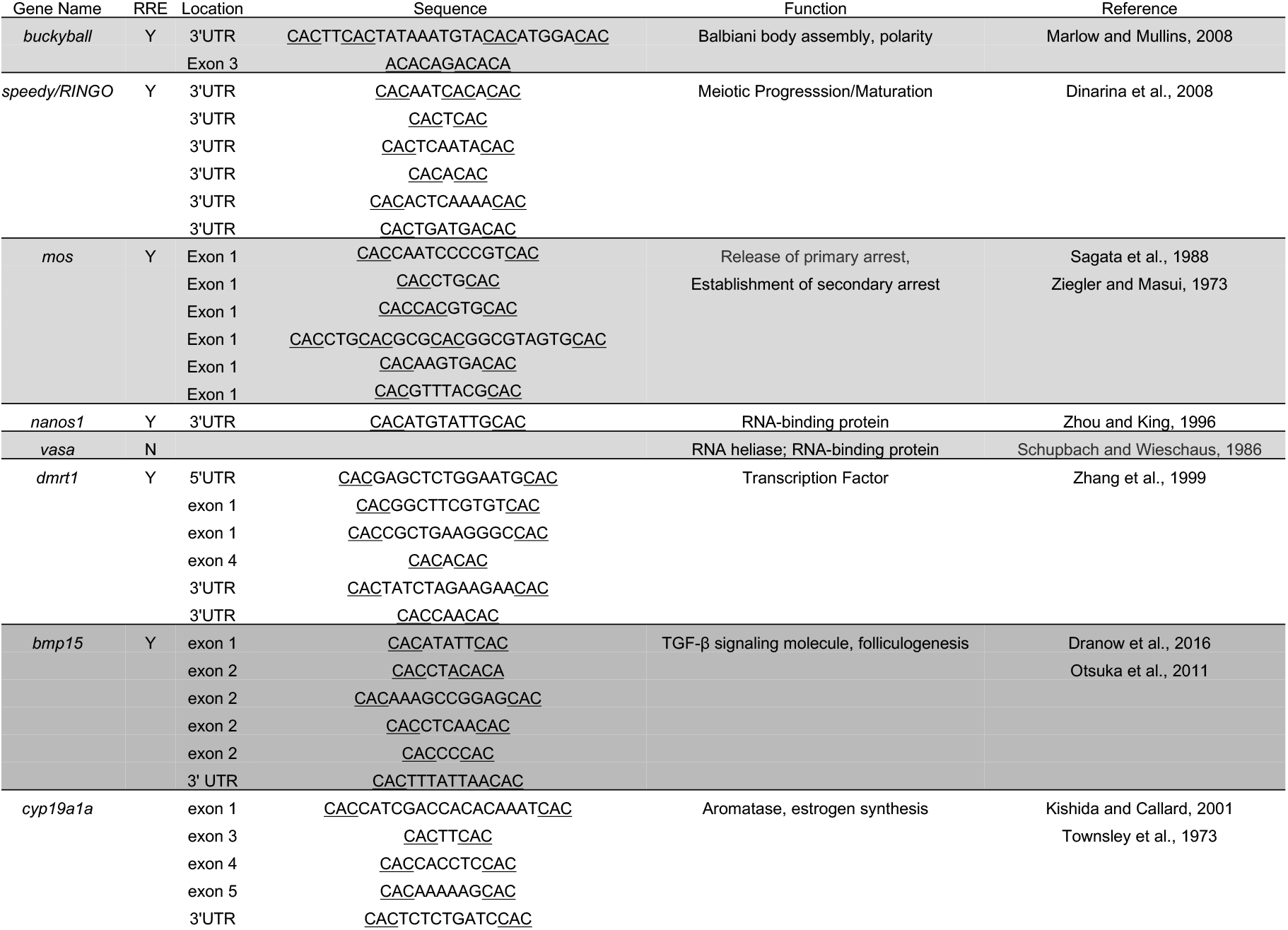
Rbpms2-binding sites in genes involved in meiosis, oogenesis, and sex determination.

### Epistasis analysis between the female factor rbpms2 and male factor dmrt1

A key and evolutionarily conserved regulator of male specific gene expression and antagonist of female fates that is necessary for male-specific development is the Double sex and Mab3 related transcription factor, Dmrt1[11, 43–50]. Mutants lacking *dmrt1* develop mostly as fertile females– the opposite phenotype of *rbpms2DMs*[11]. *rbpms2* and *dmrt1* are both expressed as RNAs in stage I oocytes (Fig. 2)[11, 27]. Because female development initiates but ultimately fails in *rbpms2DMs*, and because we identified Rbpms2 binding sites in the *3’utr* of *dmrt1* transcripts, it is possible that Rbpms2 promotes female differentiation by repressing *dmrt1* directly, and/or by antagonizing the male specific program mediated by Dmrt1. We reasoned that if *rbmps2* mutants develop as fertile males because of failed antagonism of the Dmrt1-mediated male pathway then loss of *dmrt1* should restore female development in *rbpms2DM*s. However, if *rbpms2* acts upstream of *dmrt1*, or acts in a distinct pathway then *dmrt1* mutants would fail to differentiate as females even in the absence of Rbpms2. To test this hypothesis, we conducted genetic epistasis analysis between mutants lacking *rbpms2* and *dmrt1* function. We generated adults lacking *rbpms2a/b* and *dmrt1* and screened for sex specific secondary sex traits and fertility. To determine the sex-specific identity of the germ cells in the early gonads, and to resolve the epistatic relationship between *rbpms2* and *dmrt1*, we performed immunohistochemical (IHC) analysis of d28 gonads. In triple heterozygous (TH) siblings, gonads had numerous germ cells and early-stage oocytes (Fig. 2A) or were testis-like (Fig. 2B) gonads that lacked the oocyte marker, Buc (Fig. 2B)[51, 52]. *rbpms2DM* gonads were either intersex with some early oocytes, based on presence of Buc, as well as clusters of spermatogonia-like cells (Fig. 2C), or were testis-like (Fig. 2D) as we previously reported[27]. All *dmrt1* single mutant gonads examined had early oocytes (Fig. 2E), whereas *rbpms2a;2b;dmrt1* triple mutants (TMs) (Fig. 2F) resembled their *rbpms2DM* siblings. Based on these data, *rbpms2* is required for female specific differentiation even in the absence of *dmrt1* and is therefore epistatic to *dmrt1.*

**Figure 2.**
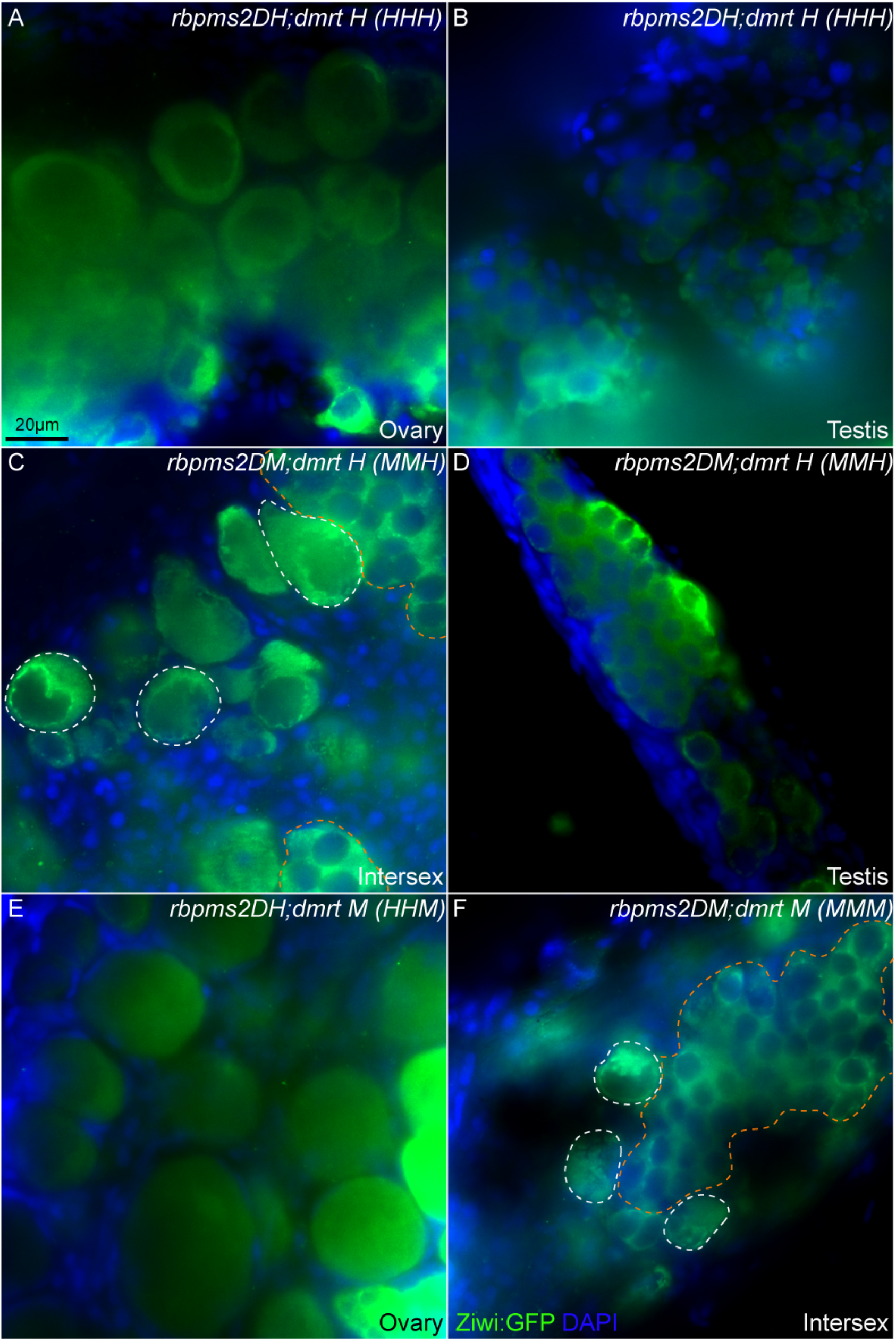
Rbpms2 is required for female gonad differentiation even in the absence of an essential male fate regulator. (A-F) Vasa and DAPI staining. Representative slice from a Z-stack reveals gonad development of d28 juvenile zebrafish, including (A) female (n=1/3) or (B) male (n=2/3) triple heterozygote *(HHH)* siblings. All *rbpms2* double mutant *(MMH)* gonads recovered were either (C) intersex (n=2/3) with oocyte-like cells outlined in white dashed lines, or testis-like cells outlined in orange dashed lines or (D) male (n=1/3). *dmrt1* mutant siblings *(HHM)* were (E) female (n=3/3), and triple mutant siblings *(MMM)* were (F) intersex (n=3/3), with oocyte-like cells outlined in white dashed lines, or testis-like cells outlined in orange dashed line. Scale bar, 20μm.

Like *rbpms2a;rbpms2bDMs* (Fig. 3A), *rbpms2;dmrt1TM* adults developed exclusively as males (Fig. 3A), indicating that *rbpms2* is also epistatic to *dmrt1* in terms of adult sex. However, unlike *rbpms2bDMs* which were fertile males, or *dmrt1* mutants with some Rbpms2 intact (compound genotypes with various *rbpms2* combinations) (Fig. 3A,B), the triple mutant males were sterile in fertility assays (failed to fertilize eggs in mating assays) (Fig. 3A,B). However, unlike the ovary or testis of “wild-type” (Fig. 3C, D), *rbpms2DM*s, or ovary of *dmrt1* mutants (Fig. 3E), TM adult males were found to lack germ cells upon inspection of the dissected gonad (Fig. 3F). It appears that germ cells of the TM gonad are lost as they attempt to transition to a male fate. Loss of the germ cells (GCs) could be due to a lack of distinct sex-specific identity (e.g. the GCs are neither truly male or female without Rbpms2 and Dmrt1) or due to a mismatch between GC and somatic cell sex-specific identities, and indicate a requirement for Rbpms2 to promote expression of factors required for female fates despite absence of *dmrt1*. Because we did not detect Rbpms2 protein in testis (Supplemental Fig.1) or in somatic cells in the ovary[27], Rbpms2 likely acts cell autonomously to promote female germline fate, which could secondarily impact somatic gonad fate. However, *dmrt1* transcripts have been detected in both germ and somatic cells[11], thus GCs may be lost in *rbpms2;dmrt1TMs* due to a cell autonomous failure to establish sex-specific identity, non-autonomous defects in somatic gonad development, or as a consequence of mismatched GC and somatic gonad identity.

**Figure 3.**
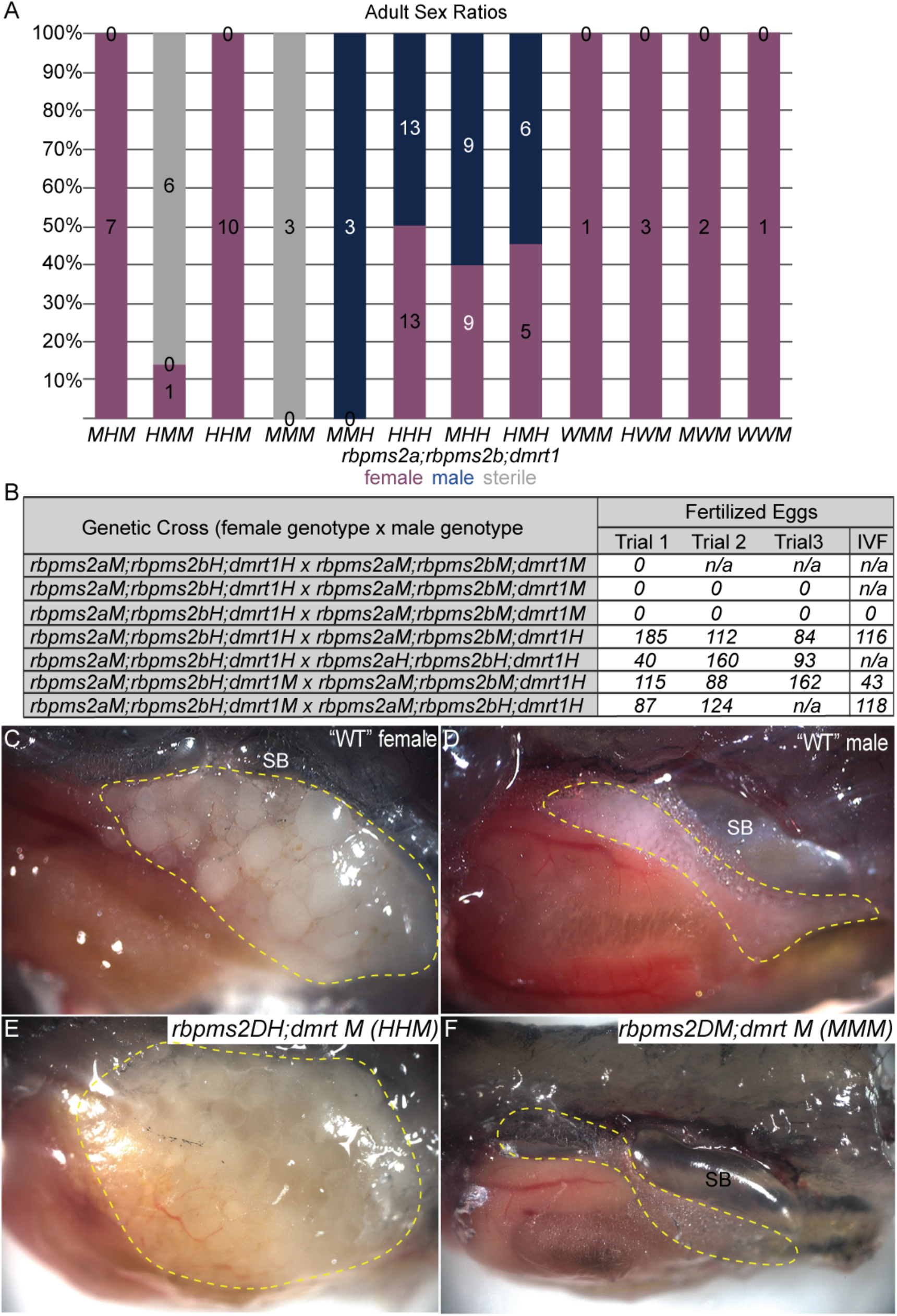
*rbpms2;dmrt1TMs* are sterile, unlike *rbpms2* double mutants. (A) Graph of adult sex ratios of progeny pooled from three triple heterozygote intercrosses for given genotypes. All double mutants also heterozygote for *dmrt (MMH)* were male as previously observed in *rbpms2* double mutants, and all *dmrt1* mutants in any other genetic combination *(HHM, MHM*, etc.) were female, as previously reported for *dmrt1*^*uc27*/uc27^, while all triple mutants *(MMM)* were sterile males. Numbers indicate number of animals for each condition. (B) Fertility was assessed in mating trials as indicated in chart. For each cross the numbers indicate number of fertilized eggs per trial for either natural breeding (eggs) or from sperm acquired through *in vitro* fertilization (IVF). n/a indicates where fish was not able to be mated for subsequent mating trials. Representative lateral view images of representative adult with tissue resected to reveal gonads (outlined in yellow dashed lines) of (C) “wild-type” female, (D) “wild-type” male, (E) *dmrti*^uc27/uc27^ female and (F) *rbpms2;dmrt1TM*. TM fish lacked morphologically obvious gonad proximal to the swim bladder (SB), compared to fertile sibling “wild-type” or single mutants.

### *Loss of dmrt1* restores bipotential and female development in the absence of *cyp19a1a*

Loss of the germ cells in *rbpms2;dmrt1TMs* adults could simply be a consequence of lack of sex-specific identity or a mismatch between GC and somatic cell sex-specific identities. If so, the gonads lacking two opposing sex-specific differentiation factors should disrupt the sex-specific identity of the gonad, potentially force a mismatch between somatic and GC identity, and recapitulate the sterile male phenotype observed in *rbpms2;dmrt1TMs*. Somatic-gonad derived factors like Dmrt1 and Cyp19a1a are continually expressed in adult gonads and loss of either gene disrupts stochastic gonad differentiation as either male or female[11, 20], with loss of *dmrt1* resulting in female sex-bias[11] and loss of *cyp19a1a* resulting in recovery of only male adults[20], indicating that the sexual phenotype of the somatic gonad must be actively maintained by the functions of these proteins. However, the genetic relationship between these two essential factors had not been appreciated in zebrafish. If bipotential gonad formation fails in *cyp19a1a* mutants due to failure to antagonize the Dmrt1-mediated male pathway, then loss of *dmrt1* should restore development of a bipotential gonad and potentially later female fates in *cyp19a1a* mutants. Conversely, if male gonadogenesis requires *dmrt1* antagonism of *cyp19a1a*, then loss of *cyp19a1a* should restore male-specific fates in *dmrt1* mutants. However, if *dmrt1* and *cyp19a1a* independently contribute to sex-specific differentiation, then *cyp19a1a;dmrt1DMs* lacking both sex-specific identity factors, would fail to confer a sex-specific identity to the somatic gonad, or result in a mismatch of soma and GC identity, leading to sterile males as observed in *rbpms2;dmrt1TMs*. To test this hypothesis, we conducted genetic epistasis analysis between mutants lacking *cyp19a1a* and *dmrt1* and determined the sex-specific identity of the double mutant germ cells using IHC analysis of d28 gonads. In *cyp19a1a*^−/+^;*dmrt1*^−/−^*(HM)* siblings, all gonads examined had numerous early-stage oocytes (Fig. 4A), consistent with the previously reported *dmrt1* mutant female-bias phenotype[11, 53]. As expected, all c*yp19a1a*^−/−^;*dmrt1*^−/+^*(MH)* sibling gonads examined were testis-like and lacked the oocyte marker, Buc (Fig. 4B)[51, 52]. In contrast, *cyp19a1a;dmrt1DM*s were either testis-like with clusters of spermatogonia-like cells that lacked Buc expression (Fig. 4C) or contained early oocytes with normal Balbiani bodies marked by Buc (Fig. 4D). Based on these data, it appears that the loss of *dmrt1* is sufficient to restore development of a bipotential gonad in the absence of *cyp19a1a*, and *dmrt1* is epistatic to *cyp19a1a* with regard to development of the bipotential gonad and initiation of female differentiation possibly by preventing *dmrt1* inhibition of *rbpms2*.

**Figure 4.**
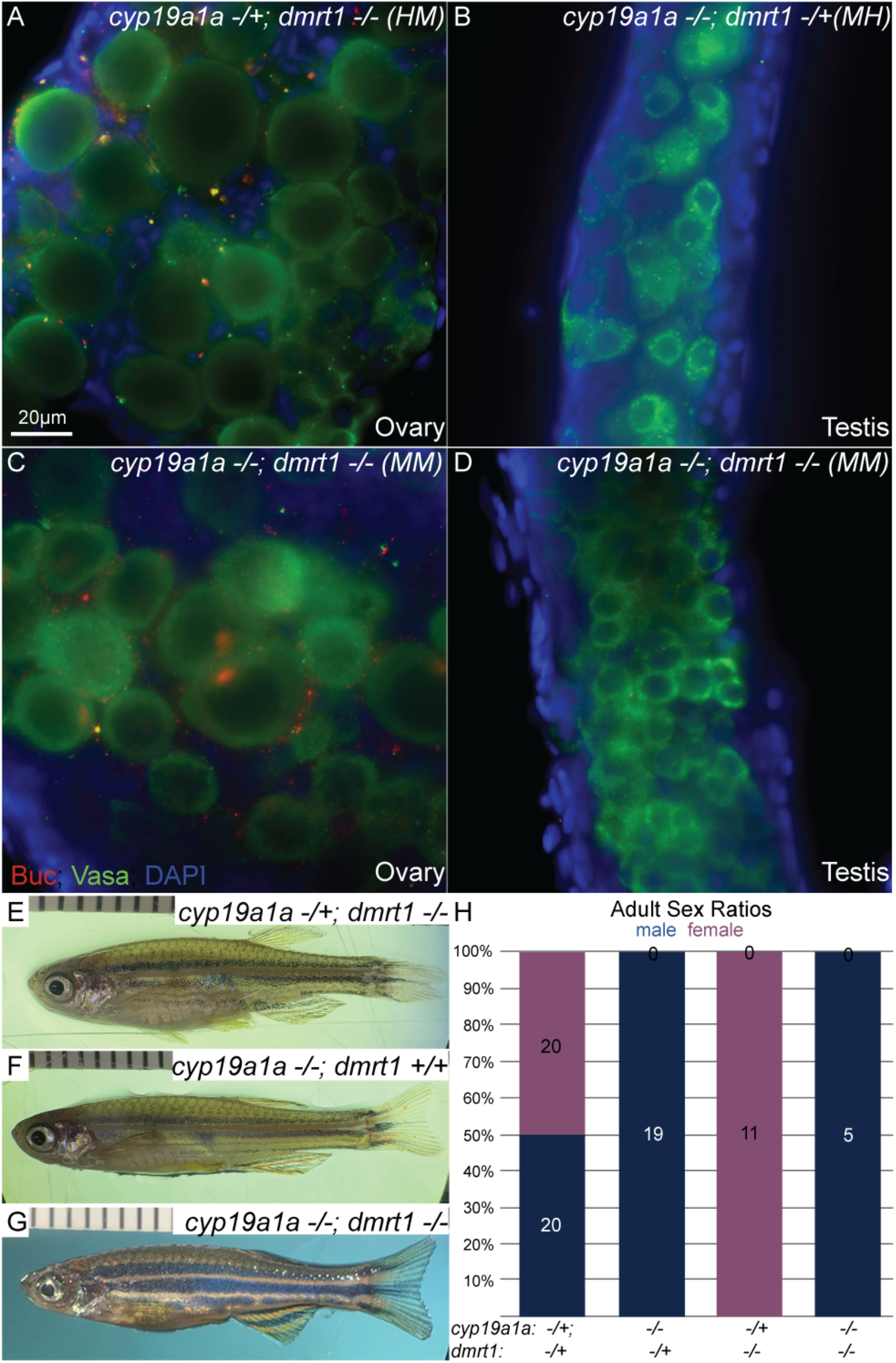
Cyp19a1a promotes bipotential and female development by inhibition of *dmrt1* and promotes later oocyte differentiation and maintenance of female fates by a Dmrt1 independent mechanism. (A-D) Vasa, Buc and DAPI staining in a representative slices from individual Z-series allows visualization of gonocyte development in bipotential gonads of d28 juvenile zebrafish, including (A) female *cyp19a1a*^*uc38/*+^; *dmrt1*^*uc27/uc27*^ *(HMs)* (n=3/3), (B) male *cyp19a1a*^*uc38/uc38*^; *dmrt1*^*uc27/+*^ *(MHs)* (n=5/5), and *cyp19a1a*^*uc38/uc38*^; *dmrt1*^*uc27/uc27*^ *(DM)* gonads that were either (C) male (n=1/4) with males germ cells or (D) female (n=3/4) with oocytes as indicated by expression of the oocyte marker, Buc (red). Scale bar, 20μm. Representative lateral view images show the external secondary sexual characteristics of d95 adults (E) *HM* female, (F) *MH* male and (G) *DM* male. (D) Graphs of adult sex ratios of progeny pooled from a heterozygote intercross and double heterozygote crossed to a mutant;heterozygote *(HH x MH)* for given genotypes. All *cyp19a1a*^*uc38*^ mutants that were also heterozygote for *dmrt1 (MH)* were males, all *dmrt1*^uc27^ mutants that were also heterozygote for *cyp19a1a* ^*uc38*^ *(HM)* were female, and all *cyp19a1a* ^*uc38/uc38*^; *dmrt1*^uc27/uc27^ double mutants were males. Numbers indicate number of animals for each condition.

To investigate the long-term effects on differentiation and fertility in the absence of *cyp19a1a* and *dmrt1*, we screened adult mutants for sex specific secondary sex traits and fertility. Despite the initial presence of early oocytes, we recovered no female *cyp19a1a;dmrt1DM* adults, indicating that oocyte differentiation eventually fails and consequently, like *cyp19a1a* mutants (Fig. 4E,H) and unlike *dmrt1* mutants (Fig. 4F,H), *cyp19a1a;dmrt1DM*s developed exclusively as males (Fig. 4G,H). Thus, Cyp19a1a function antagonizes Dmrt1 in order to establish the bipotential gonad, but also is required later in a *dmrt1*- independent manner to maintain female fate in adulthood.

Previous work showed that *cyp19a1a* is required to establish a bipotential gonad containing early oocytes[20]. In contrast to *cyp19a1a* mutants, *rbpms2 and bmp15* mutants both develop a bipotential gonad and begin to develop oocytes that ultimately fail prior to prophase I in the case of *rbpms2*[27] and after Diplotene arrest in the case of *bmp15*[20], leading to eventual differentiation of an adult testis in both cases. This suggests that Cyp19a1a is required earlier in gonad development than either Rbpms2 or Bmp15 (Fig. 5). Here we provide genetic evidence that mutation of *dmrt1* restores bipotential gonad development in the absence of *cyp19a1a* and allows for development of a gonad that contains early oocytes. That Rbpms2 is not expressed in *cyp19a1a* single mutants, which develop exclusively as males, and that normal Balbiani bodies were detected in *cyp19a1a;dmrt1DM*s indicates that inhibition of Dmrt1 allows for expression of Rbpms2 and female development in the absence of Cyp19a1a. Moreover, these results support a regulatory relationship between *dmrt1* and *cyp19a1a* and suggest that Cyp19a1a is required to establish the bipotential gonad and does so by antagonizing Dmrt1 activity (Fig. 5). However, despite the ability of *cyp19a1a;dmrt1DM*s to initially form a bipotential gonad that can develop as male or female, they fail to support ovary maintenance in adulthood, leading to the recovery of all male *cyp19a1a;dmrt1DM*s. Since the primary function of Cyp19a1a enzyme is to catalyze the final step of estrogen synthesis[18, 19], the antagonistic relationship between Cyp19a1a and *dmrt1* is likely an indirect relationship mediated by the estrogens produced by Cyp19a1a.

**Figure 5.**
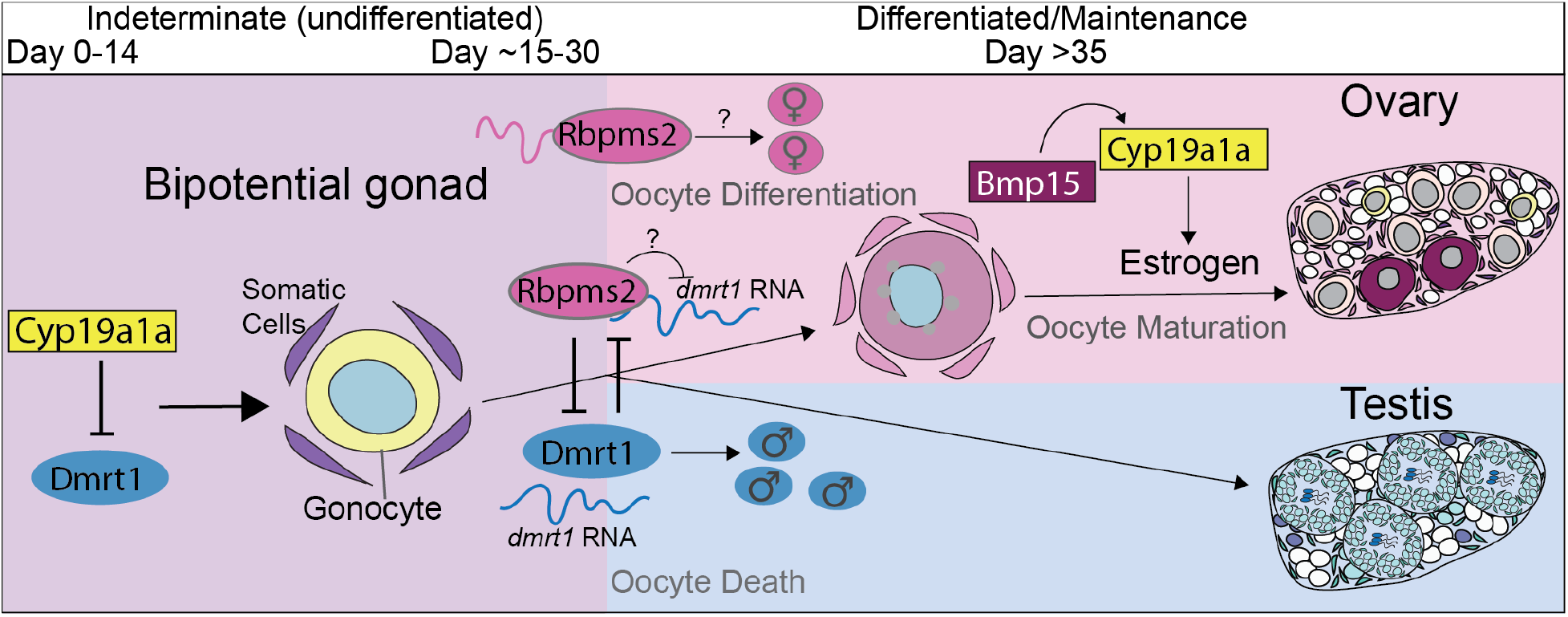
Model: *Cyp19a1a* inhibits *dmrt1* to promote bipotential gonad formation and acquisition of female fates, which requires *rbpms2* even in the absence of *dmrt1*. Timeline of requirement (from left to right) each essential regulator of sex determination shown. Cyp19a1a is required to establish the bipotential gonad by antagonizing Dmrt1 activity. Inhibition of Dmrt1 allows for expression of Rbpms2, either directly or indirectly, and female development in the absence of Cyp19a1a. Rbpms2 likely promotes female fates temporally upstream of Bmp15 and estrogen. Cyp19a1a is an enzyme that functions to catalyze the final step of estrogen synthesis; therefore, the antagonistic relationship between Cyp19a1a and Dmrt1 is likely an indirect relationship mediated by the estrogens produced by Cyp19a1a. Although Cyp19a1a is dispensable to establishment the bipotential gonad and initiation of female development in the absence of Dmrt1, it appears to be required for subsequent oocyte maturation and maintenance of female fates independent of antagonism of Dmrt1. Bmp15 produced in oocytes is required to promote oocyte differentiation and *cyp19a1a* positive granulosa cells, indicating the subsequent failure of *cyp19a1a;dmrt1DM* oocytes to mature is due to loss of the later granulosa cell-derived source of cyp19a1a,which appears not to require inhibition of Dmrt1.

Indeed, recently another group independently discovered the same epistatic relationship between *cyp19a1a* and *dmrt1* and in that work the authors provided evidence in support of estrogen involvement[53]. Additionally, that *cyp19a1a;dmrt1DMs* can establish a bipotential gonad, but eventually resolve as males, indicates that Cyp19a1a is dispensable for establishment of the bipotential gonad and initiation of female development in the absence of Dmrt1. However, this work and recent work from Wu and colleagues[53] indicate that Cyp19a1a is also required for subsequent oocyte maturation and maintenance of female fates by a mechanism that does not depend on antagonism of Dmrt1. Consistent with this notion, earlier work demonstrated that *bmp15* produced in oocytes is required to promote oocyte differentiation and *cyp19a1a* positive granulosa cells[20]. Moreover, in human ovaries and granulosa-like cells, Activin promotes Cyp19 expression via SMAD2[54, 55]; thus it seems likely that subsequent failure of *cyp19a1a;dmrt1DM* oocytes to mature is due to lack of the later granulosa cell derived source of *cyp19a1a*, which appears to utilize a Dmrt1-inhibition independent mechanism since maturation fails in the absence of Dmrt1. Finally, Rbpms2 likely promotes female fates temporally upstream of Bmp15 and estrogen since Rbpms has been previously shown to potentiate transcription mediated by Smad2/3[41], and mutants lacking estrogen[53] progress beyond diplotene stage of oogenesis in the absence of Dmrt1, whereas *rbpms2* mutants do not.

Our study identifies potential Rbpms2 targets and used genetic epistasis experiments and cell biological approaches to decipher the genetic hierarchy of critical factors involved in sex-specific differentiation of the germline and somatic gonad. We show that TGF-β signaling is activated in early GCs of the bipotential gonad and in wild-type becomes activated in the somatic gonad by d28, during the early stages of sexual differentiation. In *rbpms2DMs* initial activation of TGF-β in the germline is intact. We provide evidence that the critical female factor, *rbpms2* is epistatic to *dmrt1* in terms of adult sex. Moreover, Rbpms2 function in promoting female fates extends beyond simply repressing or antagonizing Dmrt1, as it is essential for female sex-specific differentiation even in the absence of Dmrt1. In contrast to loss of *rbpms2* function, female fates can be restored or prolonged in mutants lacking *cyp19a1a* and *dmrt1.* Taken together this work and the work of Wu and colleagues[53] indicates that *cyp19a1a* acts during at least two steps of female specific differentiation. Early *cyp19a1a-*mediated suppression of *dmrt1* is required to establish a bipotential gonad and initiate female fate acquisition in zebrafish, possibly by promoting expression of *rbpms2* which is required for female specific differentiation, even in the absence of Dmrt1. Finally, once female fates have been established Cyp19a1a is required for subsequent oocyte maturation and maintenance of female fates by a mechanism that does not depend on antagonism of Dmrt1.

## Materials and Methods

### Zebrafish

Wild-type zebrafish embryos of the AB strain were obtained from pairwise crosses and reared according to standard procedures[56]. Embryos were raised in 1X Embryo Medium at 28.5°C and staged as described[57]. All procedures and experimental protocols were in accordance with NIH guidelines and approved by ISMMS (protocol # 17–0758 INIT) IACUC. The zebrafish rbpms2a^ae30^ was previously generated in our lab using CRISPR-Cas9 mediated mutagenesis[27] and the *rbpms2b*^*sa9329*^ allele was obtained from the Sanger Institute’s Zebrafish Mutation Project[58]. The zebrafish *dmrt1*^*uc27*^, *bmp15*^*uc31*^, and *cyp19a1a*^*uc38*^ alleles were obtained from Bruce W. Draper[11, 20].

### Genotyping

#### RFLP -restriction fragment length polymorphism

Genomic DNA was extracted from adult fins, juvenile fins, or single embryos using standard procedures[56]. The genomic region surrounding *rbpms2a*^*ae30*^ was amplified using the primers 5’ - TTTGCTAAAGCCAACACGAA, 3’ -ATTCACCCTGGCCAGAGTTT, followed by digestion of the wild-type strand with the enzyme BsurI (New England Biolabs, R0581S)[27]. The genomic region surrounding *rbpms2b*^*sa9329*^ was amplified using the dCAPs primers 5’- 5’ - CACTTATCAAGCTAACTTCAAAGCAGA, 3’ -TGAAAGGGGACAAATAAGTCA, followed by digestion of the mutant strand with the enzyme MboII (New England Biolabds, R0148S) as in[27]. The genomic region surrounding *bmp15*^*uc31*^ was amplified using the primers 5’ - AGCCTTTCAGGTGGCACTCG, 3’ -GGGAGAAAGTGTTTTTCAGTGG, and *dmrt1*^*uc27*^ using primers 5’ -GTTGTAACTGGCAGCTGGAGA, 3’ -GGCGATGAGTCTGCATTTCT, but these products were resolved without digestion. After 40 cycles of PCR at 60°C annealing, samples were digested for 30 minutes using specified restriction enzyme. Digested PCR products were resolved using a 1.5% Ultrapure agarose (Invitrogen) and 1.5% Metaphor agarose (Lonza) gel.

#### HRMA-high resolution melt curve analysis

PCR and melting curve analysis were performed as described[59]. PCR reactions contained 1 μl of LC Green Plus Melting Dye (BioFire Diagnostics), 1 μl of Taq Buffer, 0.8 μl of dNTP Mixture (2.5 mM each), 1 μl of each primer (as indicated below) (5 μM), 0.05 μl of Taq (Genscript), 1 μl of genomic DNA, and water up to 10 μl. PCR and melt curve analysis were performed in a Bio-Rad CFX96 Real-Time System, using black/white 96 well plates (Bio-Rad HSP9665). PCR reaction protocol was 98°C for 1 min, then 34 cycles of 98°C for 10 sec, 60°C for 20 sec, and 72°C for 20 sec, followed by 72°C for 1 min. After the final step, the plate was heated to 95°C for 20 sec and then cooled to 4°C. Melt curve analysis was performed over a 72–92°C range and analyzed with Bio-Rad CFX Manager 3.1 software. All mutations were confirmed by TA cloning and sequencing. HRM primers as indicated; *rbpms2a*^*ae30*^ (5’ -ACACGAAGATGGCGAAGAGT, 3’ - CAGGGTGCAGGTTGGAAG), *rbpms2b*^*sa9329*^ (5’ -ATGAGGGTTCACTTATCAAGCTA, 3’ TCCGGTCAGCTGTAATGTCTAA), *dmrt1*^*uc27*^ (5’ -CTCTCGCTGCAGAAACCAC, 3’ - GGCGATGAGTCTGCATTTCT), *cyp19a1a*^*uc38*^ (5’ -CCAACTGACCTGGAATGTGTG, 3’ - AGCTACAATACTGCTGCTGCTA)

### Immunofluorescence (IF)

For whole-mount IF of gonads, trunks or tissue at d28, d45, and d120 were fixed in 4% paraformaldehyde overnight at 4°C, dehydrated in MeOH, and placed at−20°C. In the case of trunks, an incision was made to open the body cavity during staining and gonads were dissected for imaging after. Rabbit anti-Bucky ball antibody y1165 was used at 1:500 (Heim 2014). Chicken anti-Vasa antibody was a gift from Bruce W. Draper (Blokhina et al., 2019) and used at 1:3000 dilution. Alexafluor488, CY3 (Molecular Probes) secondary antibodies were diluted at 1:500. Images were acquired using a Zeiss Axio Observer inverted microscope equipped with ApotomeII and a CCD camera. Images were processed in ImageJ/FIJI, Adobe Photoshop and Adobe Illustrator.

## Author Contributions

Experiments were conceived and designed by FLM with contributions from all authors at various stages. OHK conducted RT-PCR analyses and performed pSMAD experiments. OHK, SB, and FLM generated *rbpms2;dmrtTM* lines and SR performed dissections, immunohistological, breeding and other triple mutant analyses. SR performed all other experiments and analysis in consultation with FLM. FLM contributed reagents, materials, and analysis tools. All authors contributed to various aspects of data interpretation and discussion/editing of the manuscript. SR and FLM wrote the manuscript.

## Acknowledgements

We thank members of the Marlow lab for helpful discussions, our animal-care staff for fish care (Einstein and CCMS at ISMMS) and the Microscopy CoRE at Icahn School of Mount Sinai and at Einstein. We thank Daniel Dellal and Paloma Bravo for technical assistance. Work in the Marlow lab is supported by National Institutes of Health (NIH) Grant R01-GM089979 and start-up funds to FLM. OHK was supported by NIH T32-GM007288 and NICHD F30HD082903. SR was supported by a NYSTEM training grant C32561GG and NIH F32 1F32HD097898-01A1.

