## Supplemental Figures and Legends for "Loss of *dmrt1* restores female fates in the absence of *cyp19a1a* but not *rbpms2*"

**Supplemental Figure 1. Rbpms2 is expressed in oocytes but not somatic gonad or testis.** (A) Wild-type gonad labeled with Rbpms2 antibody and DAPI reveals Rbpms2 expression within oocytes (example outlined with yellow dashed line) but not somatic gonad cells (examples outlined in pink dashed lines) or (B) testis.

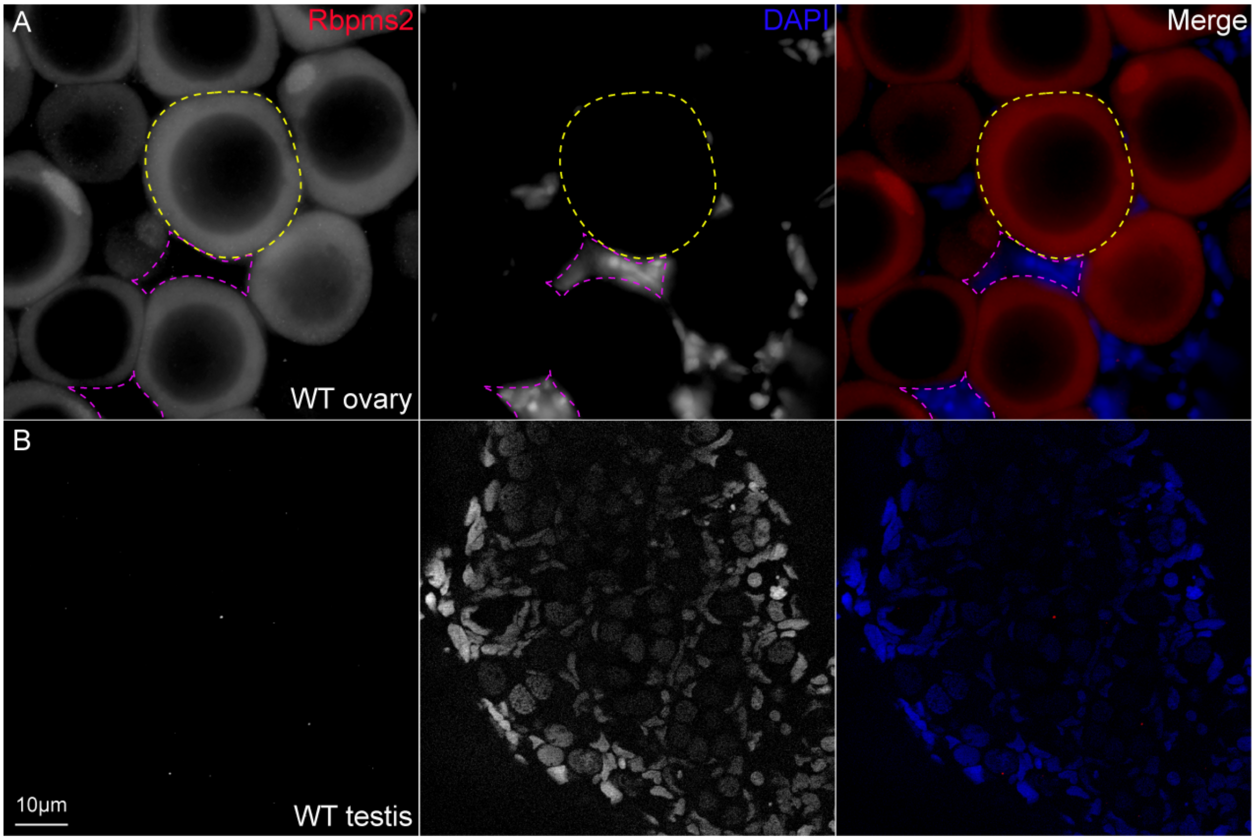
